# Sex differences in cocaine self-administration by Wistar rats after predator odor exposure

**DOI:** 10.1101/2023.02.26.530127

**Authors:** Taylor J. Templeton, Siga Diarra, Nicholas W. Gilpin

## Abstract

Traumatic stress disorders are defined in part by persistent avoidance of trauma-related contexts. Our lab uses a preclinical model of traumatic stress using predator odor (i.e., bobcat urine) in which some but not all rats exhibit persistent avoidance of odor-paired stimuli, similar to what is seen in humans. Bobcat urine exposure increases alcohol consumption in male Avoider rats, but it has not been tested for its effects on intake of other drugs. Here, we tested the effect of bobcat urine exposure on cocaine self-administration in adult male and female Wistar rats. We did not observe any effect of bobcat urine exposure on cocaine self-administration in male or female rats. We observed that (1) female rats with long access (6 hours) to cocaine self-administer more cocaine than long-access males, (2) long-access males and females exhibit escalation of cocaine intake over time, (3) stressed rats gain less weight than unstressed rats following acute predator odor exposure, (4) baseline cocaine self-administration is predictive of subsequent cocaine self-administration. The results of this study may inform future work on predator odor effects on cocaine self-administration.

## 1. Introduction

Traumatic stress disorders have a lifetime prevalence around 8% in the general population (American Psychiatric Association, 2000) with women being twice as likely as men to develop PTSD after experiencing a traumatic event (Kessler et al., 1995; Christiansen and Elklit, 2012; & Blain et al., 2010). Traumatic stress disorder symptoms are classified in four clusters that include negative affect, intrusive thoughts and memories of the traumatic event, hyperarousal, and avoidance of contexts and stimuli associated with the trauma (American Psychiatric Association, 2013). Patients with traumatic stress disorders are at higher risk for developing comorbid conditions such as substance use disorders (SUDs). Patients with SUDs, particularly cocaine use disorder (CUD), present with more severe traumatic stress disorder symptoms and respond more poorly to treatment (Najavits et al., 2007). The rate of traumatic stress disorders among cocaine users is approximately twice that of the general population (Cottler et al., 1992; Najavits et al., 1998; Back et al., 2000; Najavits et al., 1998). In patients with comorbid traumatic stress disorder and CUD, cocaine craving severity is correlated with traumatic stress disorder symptom severity (Coffey et al., 2002; Saladin et al., 2003). Currently, the specific mechanisms underlying the high prevalence of comorbid traumatic stress disorders and CUD are unclear.

Cocaine has high abuse potential. Over 1.3 million people meet criteria for CUD and over 19,000 people died from cocaine-related overdoses in 2020 (NSDUH-SAMHSA, 2020). Women exhibit higher rates of cocaine dependence (Brady et al., 1993). Although men use more cocaine than women, women experience a phenomenon in drug use known as “telescoping” in which women progress from casual use to problematic use and dependence at a faster rate than men (Brady and Randall, 1999; Kosten et al., 1993). In rat models of cocaine self-administration, females display more cocaine self-administration and drug seeking than males during acquisition, escalation, and reinstatement (Roth and Carroll, 2004). Long-access (LgA) cocaine self-administration (6-12hr/day sessions) but not short-access (ShA) (1-3hr/day sessions) produces escalation of cocaine intake (Wee et al., 2007).

Patients with a history of cocaine misuse are more likely to relapse following periods of stress, and stress potentiates cue-induced cravings in cocaine-experienced people (Preston et al., 2018). However, stress exposure does not predict the severity of cocaine cravings (Preston and Epstein, 2011). In cocaine-experienced rats, stressors (i.e., footshock, restraint, yohimbine) are highly effective at promoting reinstatement of cocaine seeking after extinction (Buffalari and See, 2009; Buffalari and See, 2011; Graf et al., 2013; McReynolds et al., 2018; Ahmed and Koob, 1997; & Erb, Shaham, and Stewart, 1996), and there are sex differences in this phenomenon. For example, female rats are more susceptible to cocaine reinstatement after yohimbine administration than males (Feltenstein et al., 2011); and relative to male rats, female rats exhibit greater reinstatement of cocaine seeking induced by the combination of a priming dose of cocaine and restraint stress (Doncheck et al., 2020).

History of cocaine use influences subsequent stress effects on cocaine seeking in rats. For example, male rats with a history of LgA self-administration (6-10hrs/day) exhibit footshock-induced reinstatement to cocaine seeking, whereas rats with ShA self-administration (1-2hrs/day) do not (Mantsch et al., 2007). Similarly, social defeat causes increases in cocaine self-administration and cocaine “binge” intake in rats (Covington and Miczek, 2001; & Quadros and Miczek, 2009). Additionally, acute- and intermittent-socially defeated male rats more readily acquire cocaine self-administration and self-administer more cocaine than non-defeated rats (Tidey and Miczek, 1997; Yap et al., 2015; & Covington et al., 2005). Interestingly, social defeat-stressed females exhibit enhanced locomotor sensitization to cocaine relative to stressed males (Holly et al., 2012).

Previous work examining predator odor stress on cocaine self-administering rats has shown that male rats that avoid Trimethylthiazoline (TMT; fox urine extract) exhibit greater sensitization to the locomotor-stimulating effects of cocaine and greater motivation to consume cocaine than non-susceptible and control rats (Brodnik et al., 2017). Another research group showed that adult male Sprague Dawley rats exposed to TMT one time exhibit increases in cocaine self-administration over time during ShA (1hr/day) and LgA (6hrs/day) cocaine sessions (Schwendt et al., 2018). Prior work examining the effects of stressful odors (e.g., feline urine, predator feces, TMT isolate) or other stressors (i.e., footshock, social defeat, restraint stress) on cocaine self-administration has been performed almost exclusively in male subjects – here, we tested predator stress effects on escalation of cocaine self-administration in male and female adult Wistar rats with long and short access to cocaine.

To do this, we used a model of traumatic stress in which rats are indexed for avoidance of a predator odor (i.e., bobcat urine)-paired context (Edwards et al. 2013; Albrechet-Souza and Gilpin, 2019). In humans, avoidance severity correlates with likelihood of diagnosis with a traumatic stress disorder, is associated with greater psychological distress after trauma, and is predictive of traumatic stress disorder symptom duration (Hendrix et al., 1994; Trappler et al., 2002; Shin et al., 2015). Furthermore, avoidance-related symptoms predict drug craving in stimulant users, suggesting that individuals with high levels of avoidance may self-medicate with psychostimulants (Somohano et al., 2019). To this point, our lab has primarily used this model to examine stress effects on alcohol self-administration in animals that do or do not exhibit persistent avoidance of predator odor-paired stimuli. In those studies, we have repeatedly observed that bobcat urine exposure produces reliable and lasting increases in alcohol self-administration in adult male Wistar rats (Edwards et al., 2013; Weera et al., 2020). Prior work from our group also shows that this procedure produces reliable increases in anxiety-like behavior (Whitaker and Gilpin, 2015) and HPA axis activation (Whitaker and Gilpin, 2015) in rats. Here, using the same model, we tested predator odor stress effects on cocaine self-administration in male and female adult Wistar rats. We hypothesized that predator odor exposure would lead to higher long-access cocaine self-administration in Avoider rats, and that this effect would be greater in females than in males.

## 2. Methods

### 2.1. Subjects

Male and female adult Wistar rats (n= 64 per sex) were obtained from Charles River Laboratories aged 8 weeks old (Raleigh, NC, USA) and were single-housed in a 12-hr reverse light cycle humidity- and temperature-controlled (22°C) colony room. Rats were allowed to acclimate to the colony room for 4 days prior to the start of surgeries. Animals were handled daily following surgery. Testing occurred during the dark period (between 8:00 am and 8:00 pm). All animals were provided with additional enrichment (Nestlet, BedderNest, and Nylabone) in their home cages throughout the study. Food was available *ad libitum* except during the first four days of the experiment while lever training. Water was always available. All procedures were approved by the Institutional Animal Care and Use Committee of the Louisiana State University Health Sciences Center and were in accordance with the National Institute of Health guidelines. For final analyses, 39 total rats were excluded due to either loss of catheter patency, post-surgical complications, or outlier behavioral responses.

### 2.2. Jugular Catheter Implantation Surgery

Rats were anesthetized with 4% isoflurane gas and maintained on 2.5% isoflurane gas throughout the surgery. They received an implanted chronic indwelling catheter into the right jugular vein using catheters made in house. The catheters were inserted under the skin directly posterior to the shoulder blades and the Silastic tubing was threaded over the right shoulder and inserted into the right jugular vein. Catheter cannulas were covered with a small piece of clean plastic tubing and a metal screw top to prevent debris from entering the catheter. Rats were allowed to recover from surgery for one hour on a heating pad before being returned to a clean home cage. 3 mL of sterile saline was administered subcutaneously to aid in recovery time. Flunixin meglumine (Covetrus; Portland, Maine, USA) was administered at the start of the surgery subcutaneously (2.5mg/kg). After the sterilized (Maxicide supplied by Henry Schein; Mandeville, LA, USA) catheter was set in place, the catheter was flushed with 0.2 mL of cefazolin/heparin (1,000 USP units per mL) solution to prevent infections and catheter clogging. Heparin was supplied by Sagent Pharmaceuticals; Schaumburg, IL, USA, and cefazolin was supplied by WG Critical Care LLC.; Paramus, New Jersey, USA. Post-operative care included weighing and visually inspecting animals daily after surgery and a catheter flush with 0.2 mL cefazolin/heparin solution once daily until the end of the experiment. Rats were allowed to recover from surgery for at least 5 days before starting experiments.

### 2.3. Drugs

Cocaine (-) HCl was generously provided by the National Institute for Drug Abuse (NIDA) Drug Supply Program and Research Triangle Institute (RTI). Cocaine was dissolved in sterile saline and delivered at a dose and volume of 0.5 mg/0.1 ml/infusion. Rats were weighed daily before the start of each session and unit cocaine dose for each rat was calculated based on body weight.

### 2.4. Drug Acquisition

To establish a baseline of cocaine intake per animal and facilitate lever-response training, rats were food restricted (20g/day for males and 15g/day for females) one day prior to the start of acquisition sessions and during the first three days of acquisition. Rats were placed into sound- and light-attenuated operant-conditioning chambers (30cm × 20cm × 24cm; Med Associates Inc., St. Albans, VT) for 2hrs/day for 10 consecutive days beginning at the same time each day. Rats began sessions 1-3 hours into the dark cycle every day for 7 days/week. Each chamber contained two retractable levers, a stimulus light above each lever, a food trough between the levers, a fan, and a speaker connected to a tone generator (ANL-926, Med Associates). A single lever press on the active lever (right lever) resulted in the delivery of a single cocaine infusion (FR1; 0.5 mg/kg/0.1 ml) along with a 20-sec presentation of 1) light above the active lever, 2) tone cue, and 3) 20-sec timeout period during which time no subsequent cocaine could occur. Presses of the left lever had no consequence but were recorded. To meet criteria for progression to the next phase of the study, rats were required to press the active lever at least 10 times during a 2-hr session for the final three consecutive days of acquisition (days 8-10) – most animals achieved this during the 10-day baseline period.

### 2.5. Stress Exposure and Conditioned Place Aversion

One day after the final day of the acquisition/baseline period, predator odor place conditioning began. No cocaine was given during this part of the experiment. All rats completed conditioned place aversion (CPA) procedure as previously described (Schreiber et al., 2017; Albrechet-Souza and Gilpin, 2019) using a 3-chamber apparatus. Briefly, on day 1 of this procedure, each rat is given five minutes to freely explore the three connected chambers in the apparatus (3-chamber pre-test), each with unique visual cues on the walls and tactile cues on the floor. To maintain an unbiased design, the chamber each rat preferred most or least relative to the other two was excluded for the remainder of the experiment. On day 2, each rat explored the two chambers it spent the most similar amounts of time in during day 1 for five minutes (2-chamber pre-test). Rats were then separated into predator odor-exposed or unstressed control groups. Again, to maintain an unbiased design, the chamber in which odor exposure would occur was assigned in a counterbalanced fashion based on chamber preference. On day 3 of the procedure, rats were confined to the “no odor” chamber in the absence of any odor for 15 min. On day 4 of the procedure, rats were confined to the “odor” chamber for 15 min, during which time rats in the odor group were exposed to bobcat urine odor and rats in the control group were exposed to no odor (same as day 3). On day 5 of the procedure, rats were once again allowed to freely move about the two chambers (2-chamber post-test). Rats were classified as “Avoiders”, or “Non-avoiders” based on the differences in time spent in the “odor” chamber between the 2-chamber pre-test and post-test. Rats with greater than a 10-sec decrease in time spent in the “odor” chamber from pre-test to post-test were categorized as “Avoiders”, and all other odor-exposed animals were categorized as “Non-avoiders”. Unstressed controls were never exposed to bobcat urine.

### 2.6. Post-stress Cocaine Self Administration

Rats were assigned to ShA or LgA self-administration groups based on their avoidance score in a counterbalanced fashion, such that Controls, Non-avoiders, and Avoiders were equally represented in the two cocaine access groups. One day following the 2-chamber post-test of the place conditioning procedure, rats were once again allowed to self-administer cocaine (0.5 mg/0.1 ml/infusion) on an FR1 schedule of reinforcement for either 1hr/day (ShA) or 6hrs/day (LgA) for 10 consecutive days in the same operant chambers used for acquisition and baseline. Specific operant boxes were used for males only, and others were used for females only during the entirety of the experiment. At the end of the study, one day after the 10^th^ day of self-administration, rats were allowed to self-administer cocaine for 1 hour and then anesthetized with 4% isoflurane gas, decapitated, and the brains were flash frozen in 2-methylbutane for future analysis.

### 2.7. Catheter Patency

Catheter patency was checked once per week by flushing catheters with 0.2 mL Brevitol Sodium (distributed by Henry Schein; Mandeville, LA, USA) and observing body response. Rats that failed to go limp after three seconds were excluded from the study.

### 2.8. Statistical Analysis

All data was analyzed using Prism Graph Pad version 9.0 (GraphPad Software, Inc., La Jolla, CA, USA). Data were analyzed using ANOVAs with stress groups, sex, and self-administration access groups as between-subjects factors and session days as a within-subjects factor when appropriate. Significant effects were followed up with Tukey’s or Bonferonni post-hoc tests where appropriate (Bonferonni post-hoc tests were used for all repeated measures ANOVAs). Outliers were detected using the interquartile range (IQR) rule (Jones, 2019), and data analyzed after outliers were detected were analyzed using Mann-Whitney test. Correlations were performed using Pearson correlations. Statistical significance was set at p<0.05.

## 3. Results

### 3.1. Avoidance of predator odor context is not associated with cocaine self-administration

Adult male and female Wistar rats were exposed to bobcat urine and indexed for avoidance behavior of the odor-paired chamber (see **Fig. 1**), and 62% of males and 46% of females were categorized as Avoiders. A Pearson correlation revealed no association between avoidance and cocaine deliveries in ShA stressed males (r=0.30; *p=*0.37), LgA stressed males (r=0.25; *p=*0.41), ShA stressed females (r=-0.05; *p=*0.89), or LgA stressed females (r=-0.07; *p=*0.83). We analyzed the effects of avoidance phenotype and time on cocaine deliveries using separate two-way repeated measures (RM) ANOVAs for LgA males, LgA females, ShA males, and ShA females. There was no effect of avoidance phenotype on post-stress cocaine deliveries in any of these groups (p>0.05 in all cases). Because we did not find differences in cocaine intake between Avoiders and Non-avoiders, we collapsed Avoiders and Non-avoiders into a single group of “Stressed” rats for each sex, and we compared Stressed rats to unstressed Controls in all subsequent analyses.

**Fig 1.**
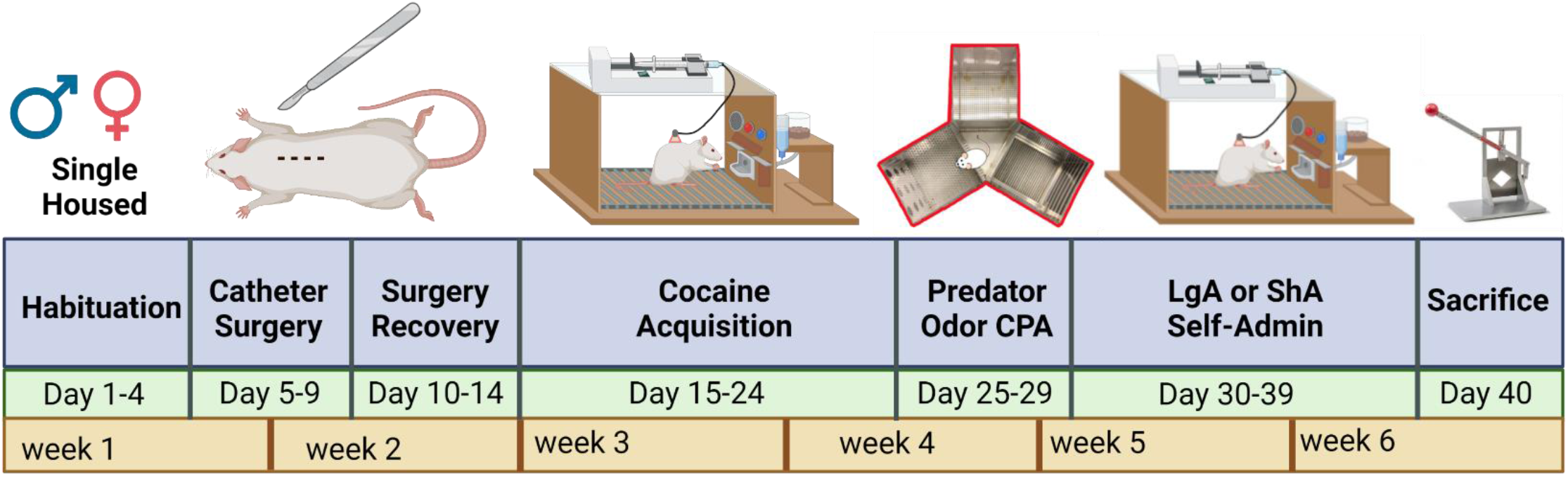
Experiment timeline. Each cohort was treated identically upon arrival to the LSUHSC animal facility. The experiment lasted 26 days from the start of cocaine exposure to sacrifice. All rats acquired cocaine operant conditioning. Then rats underwent place aversion conditioning with bobcat urine. Rats in the control group never experienced bobcat urine. Avoidance was measured and subjects were divided into ShA or LgA groups. The ShA and LgA groups completed 10 days of post-CPA self-administration.

### 3.2. Predator odor exposure does not alter cocaine self-administration

A 2-way RM ANOVA showed that LgA males escalated cocaine drug deliveries regardless of stress history (F[2.41,55.64]=6.29; p<0.01), but there was no effect of stress (F[1,23]=0.02; *p=*0.88) (**Fig. 2A**). Neither stress (F[1,20]=2.11; *p=*0.16) nor day (F[1.93,38.53]=1.92; *p=*0.16) had effects on ShA males’ cocaine deliveries (**Fig. 2A**). **Figure 2B** highlights that LgA males exhibit greater increases in cocaine deliveries over time relative to ShA males (F[1,43]=27.94; p<0.0001) in the absence of stress effects (F[1,43]=1.90; *p=*0.18).

**Fig. 2:**
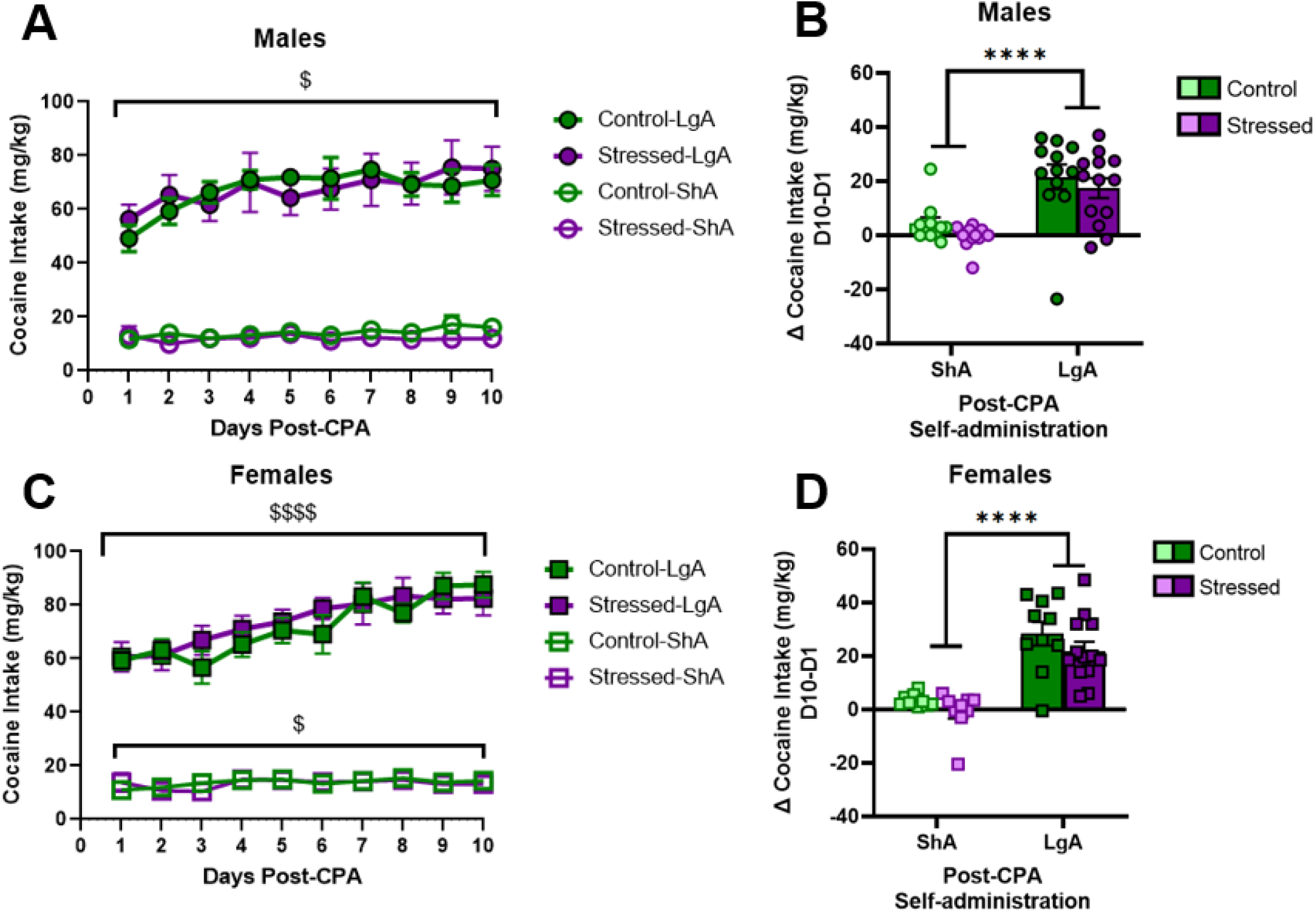
Long-access males and females escalate self-administration; no effect of stress. **A-B)** LgA males escalated cocaine intake (*p=*0.01), but ShA males did not change intake over time. **C-D)** LgA females escalated cocaine intake. Data presented as mean ± SEM. $ denotes *p*<0.05 and a main effect of time; and **** denotes *p*<0.0001 and a main effect of access condition.

A separate 2-way RM ANOVA showed that LgA females escalated cocaine deliveries regardless of stress history (F[2.15,45.22]=26.18; p<0.0001; **Fig. 2C**), and again there was no effect of stress (F[1,21]=0.21; *p=*0.65). A separate 2-way RM ANOVA in ShA females revealed a main effect of day (F[2.48,42.13]=3.29; *p=*0.04) but not of stress (F[1,17]=0.05; *p=*0.83) (**Fig. 2C**). **Figure 2D** highlights LgA females exhibit greater increases in cocaine intake over time relative to ShA females (F[1,38]=54.74; p<0.0001) in the absence of stress effects (F[1,38]=2.738; *p=*0.11).

### 3.3. Males and females escalate cocaine deliveries

In separate analyses, we compared self-administration across sexes in animals with the same access conditions (LgA or ShA) collapsed across stress history. In a 2-way RM ANOVA analysis of cocaine delivery data in LgA animals, we observed that all rats escalate cocaine deliveries over 10 days (F[2.50,115.0]=22.93; p<0.0001; **Fig. 3A**). There was no main effect of sex (F[1,46]=3.35; *p=*0.07; **Fig. 3A**). ShA males and females did not differ in intake (F[1,39]=0.23; *p=*0.64), but there was a main effect of day (F[2.46,95.82]=3.81; *p=*0.02) on ShA cocaine delivery showing all rats escalated cocaine deliveries over time. A 2-way ANOVA on cocaine deliveries in LgA rats showed LgA males and females take more cocaine during the second half of post-CPA self-administration (days 6-10) than during days 1-5 (D1-5) of post-CPA self-administration (main effect of access days: F[1,46]=21.52; p<0.0001; **Fig. 3B**). In LgA rats, females take more cocaine than males during self-administration days 6-10 (D6-10) (days x sex interaction: F[1,46]=2.750; p<0.01; **Fig. 3B**).

**Fig. 3:**
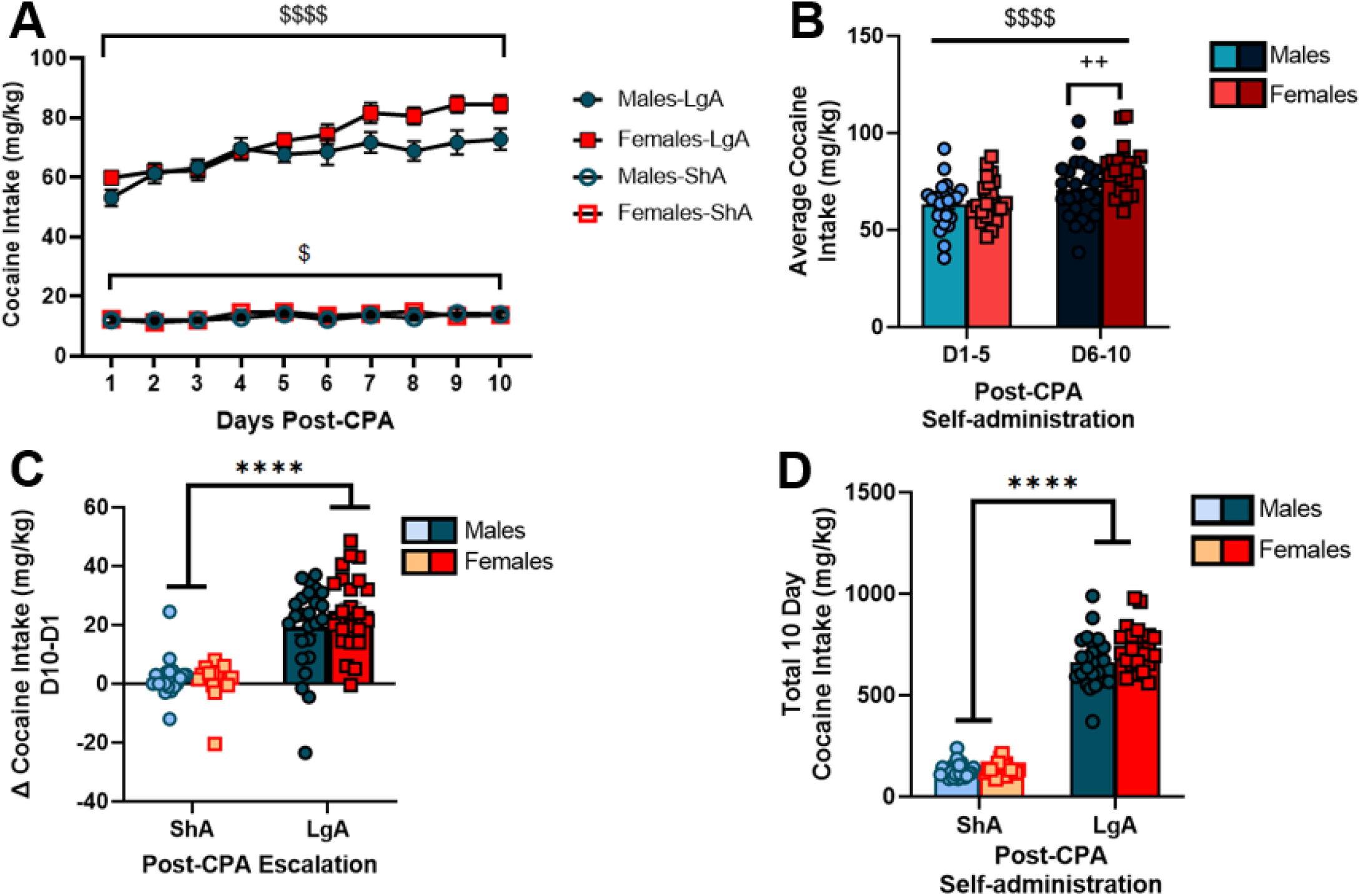
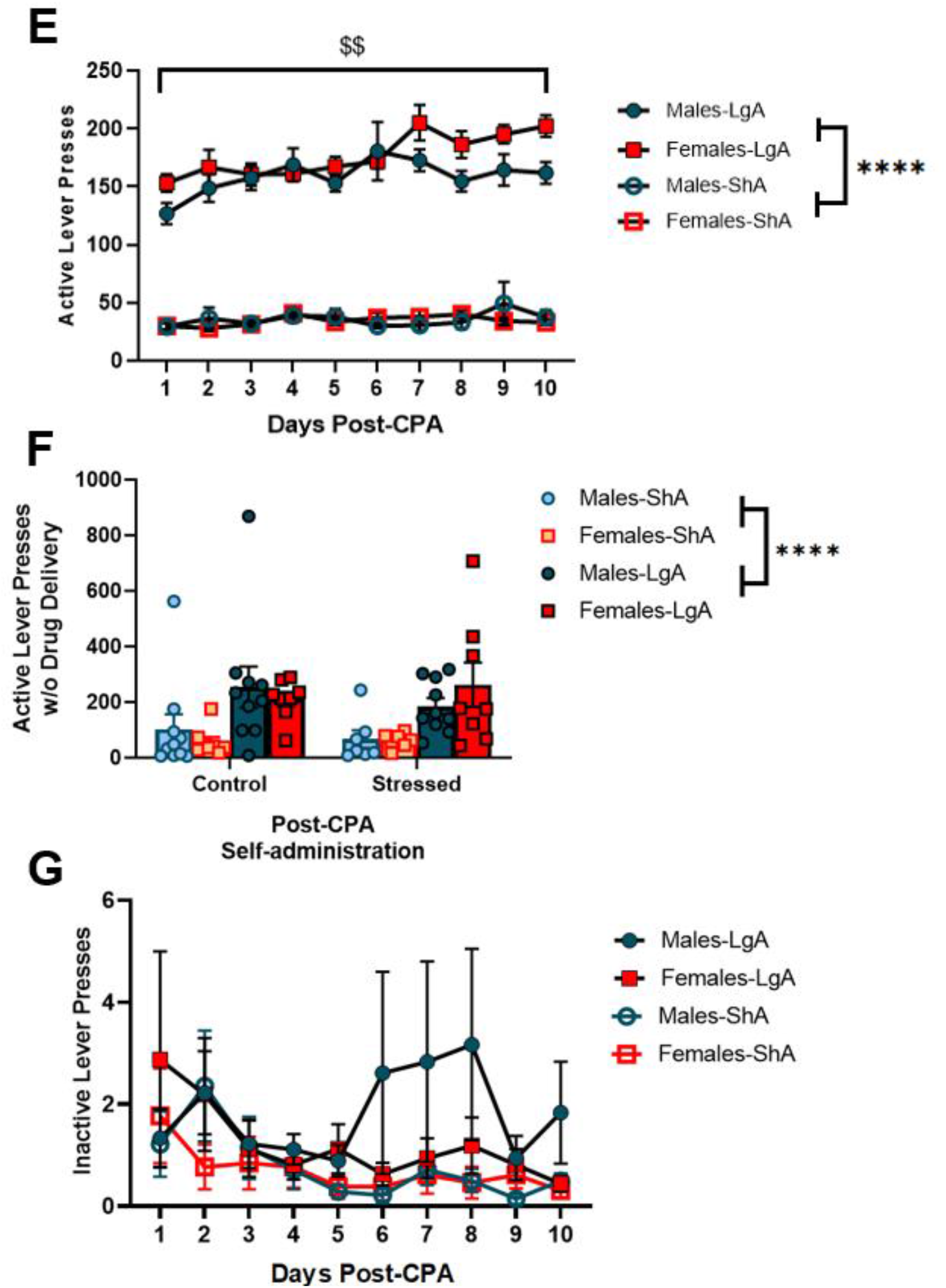
LgA females self-administer more cocaine than LgA males on days 6-10. **A)** All LgA males and females (combined control and stressed groups) escalated cocaine intake over 10 days post-CPA self-administration. **B)** LgA males and females received more cocaine deliveries during the second half of post-CPA self-administration, and females received more cocaine deliveries than males on days 6-10. **C)** LgA males and females escalated cocaine intake, but ShA rats maintained similar intake levels throughout the 10-day testing period. LgA males and females escalated intake similar amounts across all 10 days. **D)** All LgA rats received more cocaine deliveries than ShA rats. **E)** LgA males and females increased total active lever presses (with and without drug delivery) over time. **F)** LgA males and females pressed the active lever more during 20-sec time-outs than ShA males and females; there was no effect of stress. **G)** There was no difference between how many inactive lever presses males and females made during post-CPA self-administration sessions. * denotes main effect of access condition, $ denotes main effect of day, # denotes main effect of sex, and + denotes interaction effects. **** denotes *p*<0.0001; *** denotes *p*<0.001; ** denotes *p*<0.01; * denotes *p*<0.05.

A follow-up 2-way ANOVA highlights that LgA rats display greater escalation of cocaine deliveries than ShA rats (F[1,85]=76.46; p<0.0001; **Fig. 3C**). LgA males and females escalated drug deliveries similar amounts across all 10 days post-CPA self-administration (F[1,85]=1.06; *p=*0.31). When comparing total cocaine deliveries, we found all LgA rats received more cocaine deliveries than ShA rats (F[1,85]=876.9; p<0.0001; **Fig. 3D**). There was not a significant difference between sexes in total deliveries in 10 days, but there was a trend that showed females self-administered more than males (F[1,85]=3.53; *p=*0.06; **Fig. 3D**). **Figure 3E** illustrates LgA males and females increase total active lever presses (with and without drug delivery) over time (F[3.20,107.5]=4.21; p<0.01) with no sex differences (F[1,34]=2.96; *p=*0.09). There was no change in active lever presses across days (F[1.81,51.56]=1.07; *p=*0.34) nor sex (F[1,30]=0.04; *p=*0.84) in ShA rats (**Fig. 3E**). A 3-way ANOVA on total active lever presses during the time-out period, when no cocaine was available, revealed that LgA rats press the active lever more during 20-s timeouts than ShA rats (F[1,60]=18.70; p<0.001; **Fig. 3F**); there was no effect of stress (F[1,60]=0.11; *p=*0.74) nor sex (F[1,60]=0.013; *p=*0.91) on responding during time-out periods.

**Figure 3G** shows inactive lever presses (lever presses resulting in no reinforcer delivery) for all LgA and ShA males and females during post-CPA sessions. We used a 2-way RM ANOVAs to analyze sex and time effects on inactive lever presses in LgA and ShA animals. After removing outliers (1 male) identified using the IQR test, a Mann-Whitney test revealed that there were no differences in inactive lever pressing during post-CPA self-administration between any group. There was no time x sex interactions between LgA males and LgA females (F[9, 288]=0.82; *p=*0.60), and no time x sex interactions between ShA males and females F[9, 225]=0.87; *p=*0.56). There was no difference in inactive lever presses between ShA and LgA animals (F[27, 513]=0.65; *p=*0.91). Outlier data appeared to be associated with traditional stimulant-induced stereotypy behavior.

### 3.4. Baseline cocaine intake correlates with post-CPA cocaine self-administration

Collapsing across stress groups, in the ShA male, ShA female, and LgA female groups, baseline cocaine intake (average cocaine intake [mg/kg] during acquisition days 8, 9, and 10) was positively correlated with post-CPA cocaine intake (ShA males [r=0.66; p<0.001; **Fig. 4A**]; ShA females [r=0.57; *p=*0.01; **Fig. 4C**]; LgA females [r=0.50; *p=*0.01; **Fig. 4D**]. LgA males exhibited a trend toward the same correlation (r=0.40; *p=*0.058; **Fig. 4B**).

**Fig. 4:**
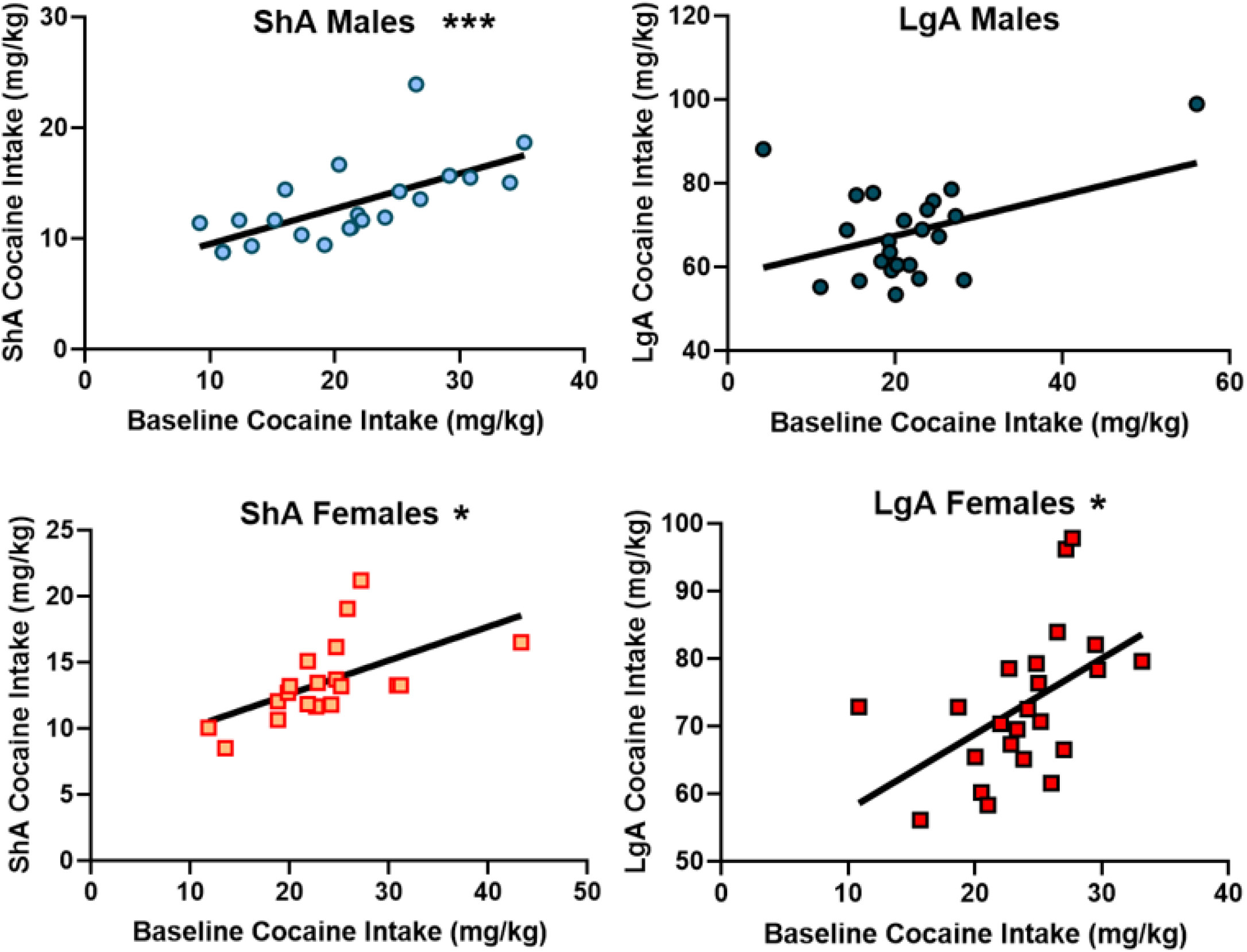
Baseline cocaine intake predicts future cocaine intake. **A-D)** Male and female ShA and LgA rats that were low responders during acquisition continued to be low responders and high responders continued to be high responders after the conditioned place aversion test regardless of stress exposure group and access group. *denotes *p*<0.05.

### 3.5. Weight gain during self-administration and stress exposure

Rats were weighed daily throughout the experiment. We analyzed percent change in body weight during the pre-CPA cocaine acquisition period (2hrs/day cocaine access), and during the post-CPA LgA (6hrs/day cocaine access) and ShA (1hr/day cocaine access) period. Males and females increase body weight similarly during acquisition (unpaired t-test: *p=*0.59; **Fig. 5A**). During post-stress self-administration, all (control and stressed combined) LgA males gained less weight than all ShA males (F[1,85]=7.04; p<0.01); and there was a sex x access condition interaction effect on body weight gain (F[1,85)=8.38; p<0.01; **Fig. 5B**).

**Fig. 5:**
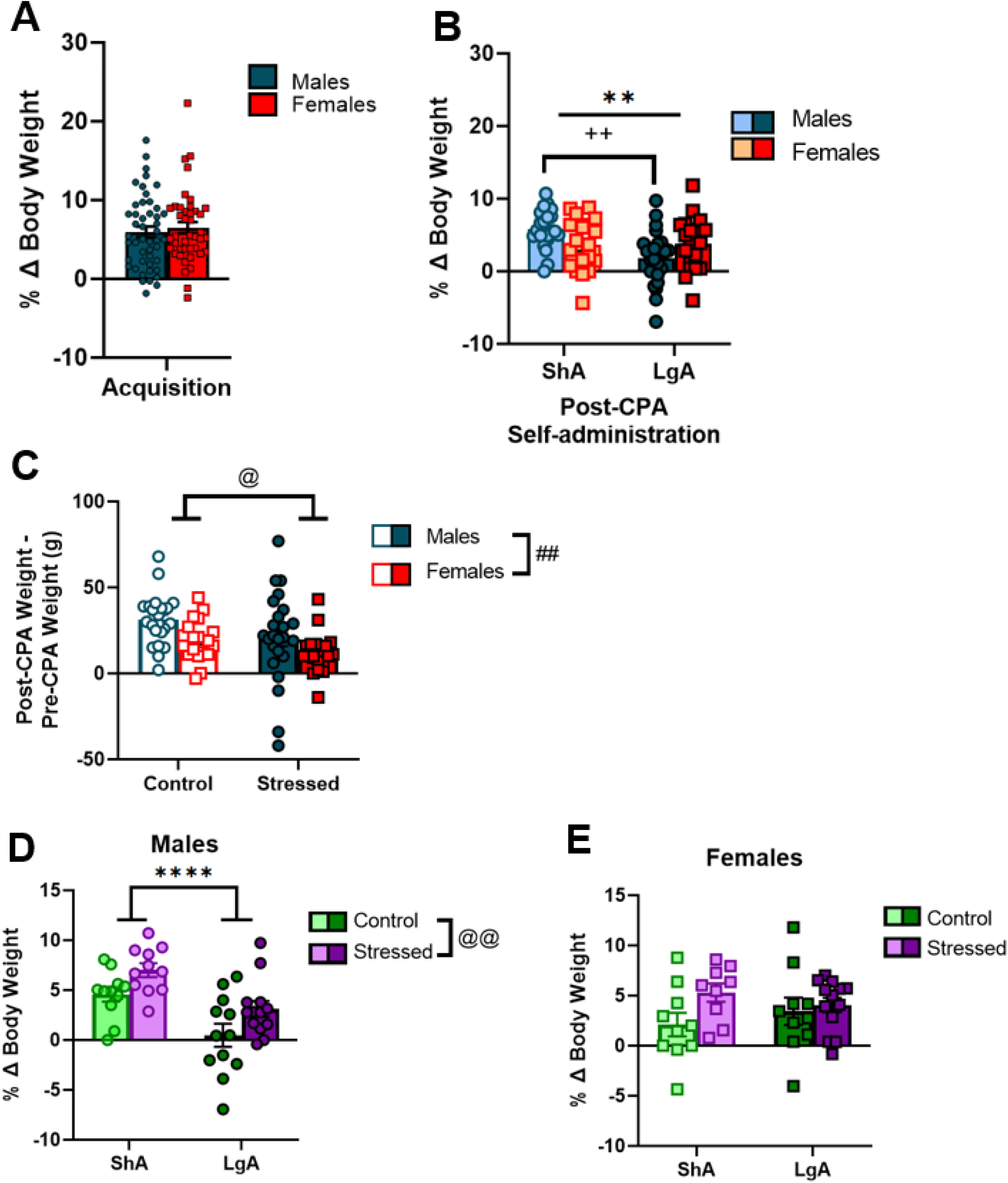
Weight gain during acquisition, CPA, and post-CPA ShA and LgA. **A)** Males and females increased body weight similarly during acquisition. **B)** LgA males gained less weight than ShA males while females were unaffected. **C)** Stressed males and females gained less weight than controls following acute predator odor exposure across the 5-day CPA procedure. **D)** LgA males gained less weight than ShA males, and stressed males gained more weight than control males after predator odor exposure during post-CPA 10-day self-administration. **E)** There was no difference in post-CPA weight gain between LgA and ShA females nor control and stressed females during 10-days post-CPA self-administration. * denotes main effect of access condition, $ denotes main effect of day, # denotes main effect of sex, @ denotes main effect of stress, and + denotes interaction effects. **** denotes *p*<0.0001; *** denotes *p*<0.001; ** denotes *p*<0.01; * denotes *p*<0.05.

Next, we collapsed data across LgA and ShA conditions to examine acute effects of predator odor exposure (i.e., during the 5-day conditioning procedure) on body weight gain in male and female rats. When comparing body weight (g) 48hrs after the CPA test (i.e., day 1 of post-CPA self-administration prior to the start of the operant session) to body weight prior to day 1 of the conditioning procedure, we observed that stressed males and females gain less weight than controls (F[1,85]=5.59; *p=*0.02; **Fig. 5C**), and females gained less weight overall than males (F[1,85]=8.16; p<0.01; **Fig. 5C**) during the 5 days of the conditioning procedure, presumably due to acute stress effects of the predator odor exposure.

Finally, we used a 2-way ANOVA to analyze percent body weight gain in male rats during the 10 days of post-CPA self-administration. LgA males gain less weight than ShA males (F[1,43]=20.85; p<0.0001; **Fig. 5D**), and stressed males gain more weight than control males during the 10 days of post-CPA self-administration (F[1,43]=8.41; p<0.01; **Fig. 5D**). A separate 2-way ANOVA of percent body weight gain in female rats during the 10 days of post-CPA self-administration revealed there was no difference in post-CPA body weight gain between LgA and ShA females (F[1,38]=0.002; *p=*0.97), nor was there a stress effect (F[1,38]=3.28; *p=*0.08; **Fig. 5E**) on body weight gain in females during the 10-day post-CPA self-administration period.

## 4. Discussion

The main results of this experiment demonstrate that 1) acute predator odor stress does not affect ShA or LgA cocaine self-administration in adult male and female Wistar rats; 2) male and female rats exhibit greatest escalation of cocaine intake under long-access conditions; 3) females self-administer more cocaine than males; and 4) early cocaine intake during 2-hr acquisition periods is positively associated with subsequent levels of cocaine intake in males and females under LgA and ShA conditions. We also report that 1) stressed rats gain less weight in the days immediately following bobcat urine exposure than unstressed controls; 2) LgA males gain less weight than ShA males, and surprisingly, 3) stressed males (regardless of access condition) gain more weight than unstressed control males during 10 days of post-CPA self-administration. Interestingly, cocaine access condition and stress exposure did not affect body weight gain in females.

In this study, females acquired cocaine at a similar rate to males; then they self-administered more under long-access conditions, but this was only seen in the later session (LgA D6-10). We originally hypothesized that Avoider rats would take more cocaine than Non-avoiders and unstressed rats and that females would take more than males. Our data indicated that Avoiders did not differ in their cocaine intake from Non-avoiders nor unstressed rats. Furthermore, all stressed rats (Avoiders + Non-avoiders) did not self-administer more cocaine than unstressed rats. This differs from previous results from our lab in which bobcat urine exposure produces long lasting increases in alcohol self-administration in Avoiders (Edwards et al., 2013; Weera et al., 2020) and blunts alcohol aversion (Schreiber et al., 2019). It is possible that a single odor exposure was not sufficient to change cocaine self-administration due to the strong reinforcing properties of cocaine.

Prior work from another research group also reported that a one-time exposure to predator odor (TMT in this case) failed to alter LgA cocaine self-administration (FR1; 0.33mg/infusion) (Schwendt et al., 2018). That work differed from ours in that their animals were exposed to predator odor without prior cocaine experience, subjects were Sprague Dawley rats, and LgA sessions began 23 days after a ten-minute TMT exposure. That prior study reported that TMT-stress resilient rats, identified according to anxiety-like behavior and startle reactivity, self-administered fewer cocaine infusions during ShA sessions compared to controls and TMT-stress susceptible rats. Comparing between experiments, Sprague-Dawley rats in the Schwendt et al. (2018) study responded less for cocaine and received fewer cocaine infusions during ShA (1hr/day) and LgA (6hr/day) sessions than Wistar rats in the current study (2-to-4 fold higher responding in the current study). These differences may be attributable to strain and/or procedural differences between the studies (timing of self-administration acquisition, type of odor, etc.). Future work may benefit from utilizing repeated stress exposures and/or fewer baseline acquisition sessions prior to odor exposure.

Bobcat urine is an innately stressful stimulus that triggers the release of pro-stress hormones (e.g., corticosterone) and increases anxiety-like behaviors in rats (Whitaker & Gilpin, 2015; Kondoh et al., 2016; Takahashi et al., 2005; Albrechet-Souza et al., 2020; & Hegab and Wei, 2014). A subset of rats exposed to bobcat urine exhibit long-lasting and extinction-resistant avoidance of stimuli paired with that odor (Edwards, et al., 2013). Other work reported that cat urine-exposed rats exhibit less habituation in the acoustic startle response than unstressed controls (Hadad et al., 2016). Interestingly, female rats exposed to bobcat urine exhibit lower acoustic startle reactivity relative to unstressed control females, unlike what is seen in males, highlighting sex differences in the behavioral response to bobcat urine exposure in male and female rats (Albrechet-Souza et al., 2020).

Previous work in our lab reported that bobcat urine exposure reduces body weight gain in Avoiders and Non-avoiders relative to unstressed controls (Albrechet-Souza et al., 2020; Weera et al., 2020). Here, we confirmed those findings such that stressed animals gained less weight during the week in which bobcat urine exposure and place conditioning occurred. Interestingly, however, predator odor-exposed males subsequently gained more weight than unstressed controls during the following 10 days of cocaine self-administration, regardless of access condition. Cocaine is frequently used as a weight control strategy, more so in women than in men (Cochrane & Brewerton, 1998; Bruening et al., 2018). One study reported that cocaine-dependent men have lower body mass index than non-dependent men, but report more compulsive eating and consume more calorie-dense food indicating cocaine’s weight loss effects may be due to changes in nutrient metabolism rather than changes in feeding behavior (felt, et al., 2013).

Preclinical work has reported that cocaine does not change long-term food intake in rats, but rather cocaine has acute anorexigenic effects (Balopole, et al., 1979). It has been suggested that increases in mesolimbic dopamine may suppress food consumption, similar to what is seen following administration of the dopamine type-2 receptor agonist, bromocriptine (Thanos et al., 2011). However, following acute cocaine-induced appetite suppression, rats display increased motivation for high-calorie food which may lead to “binge eating” in cocaine-experienced rats (Laque et al., 2022). Clinical and preclinical work suggests that subjects with a history of cocaine use consume more calories and more high-fat food, but they do not gain more weight than non-cocaine users (Ersche et al., 2013; Bane et al., 1993). It has been speculated that these effects may be due to cocaine stimulation of cocaine and amphetamine-regulated transcript (CART) release and activation of CART-expressing nucleus accumbens neurons, as well as subsequent reductions in leptin production (Wortley et al., 2004; Vicentic and Jones, 2007; & Rogge et al., 2008). Furthermore, cocaine’s ability to antagonize the serotonin transporter and norepinephrine transporter (resulting in a decrease in pre-synaptic serotonin and norepinephrine reuptake) likely result in increased appetite, decreased leptin production, and increased thermogenesis (Yadav et al., 2009 & Rayner and Trayhurn, 2001). Here, we report that males with extended access (6hrs/day) to cocaine exhibit attenuated weight gain across 10 days of long-access cocaine self-administration relative to males with shorter access (1hr/day) to cocaine. One limitation of this study is that we did not measure food intake over time.

Intravenous self-administration experiments report that female rats respond more for cocaine than males at all stages of access (i.e., acquisition, escalation) (Roth & Carroll, 2004; Lynch & Carroll, 1999). Female rodents also more readily reinstate drug use after a period of abstinence in the absence of any reinforcing cues when compared to males (Anker & Carroll, 2010a; Buffalari et al., 2012). Here, in adult male and female Wistar rats, we confirmed prior work showing that LgA cocaine self-administration but not ShA produces escalation of cocaine intake (Wee et al., 2007). Women begin using cocaine earlier in life (Griffin et al., 1989), develop CUD faster than men (SAMHSA NSDUH 2016; Brady et al., 1993; Quiñones-Jenab & Jenab, 2012), experience more cocaine-related stress, paranoia, and hyperalgesia than men (Back et. al., 2005), and seek treatment for CUD more times in their lives than men (Becker and Hu, 2008). It is possible that different strategies will be effective at reducing cocaine use in men versus women. Furthermore, sex/gender differences in cocaine reward may depend on menstrual cycles and circulating/brain estradiol and progesterone in women (Evans et al., 2002; Evans and Foltin, 2006; Reed and Carr, 2000; Moran-Santa Maria et al., 2018; Sinha et al., 2007; Fox et al., 2013; Milivojevic et al., 2016; Sofuoglu et al., 2004). Unfortunately, we did not monitor the estrous cycles of females, which is a limitation of this study considering that female rodents self-administer more cocaine during estrus/proestrus than diestrus and when given estradiol (Lacy, Austin, and Strickland, 2019; Jackson et al., 2006).

In conclusion, this study confirms various prior findings in the cocaine self-administration and predator odor stress fields. Prior findings showing that bobcat urine exposure produces escalation of alcohol self-administration did not generalize to cocaine self-administration in this study regardless of sex and cocaine access condition. Strategies for future work in this area may consider exposing animals to repeated stress, changing the timing of stress relative to cocaine self-administration, or exploring other predator odors.

## Acknowledgments

We thank Dr. Mary Secci, Chuck Jager, and Kim Edwards for their technical assistance.

## REFERENCES

Ahmed, S. H., & Koob, G. F. (1997). Cocaine-but not food-seeking behavior is reinstated by stress after extinction. Psychopharmacology, 132(3), 289–295. https://doi.org/10.1007/s002130050347

Albrechet-Souza, L., & Gilpin, N. W. (2019). The predator odor avoidance model of post-traumatic stress disorder in rats. Behavioural Pharmacology, 30(2 and 3-Spec Issue), 105–114. https://doi.org/10.1097/FBP.0000000000000460

Albrechet-Souza, L., Schratz, C. L., & Gilpin, N. W. (2020). Sex differences in traumatic stress reactivity in rats with and without a history of alcohol drinking. Biology of Sex Differences, 11(1), 27. https://doi.org/10.1186/s13293-020-00303-w

American Psychiatric Association. (2000). Diagnostic and Statistical Manual of Mental Disorders: (DSM-4. 4th ed.). https://dsm.psychiatryonline.org/doi/epdf/10.1176/appi.books.9780890420249.dsm-iv-tr

American Psychiatric Association. (2013). Diagnostic and Statistical Manual of Mental Disorders: (DSM-5. 5th ed.). https://dsm.psychiatryonline.org/doi/epdf/10.1176/appi.books.9780890425596

Anker, J. J., & Carroll, M. E. (2010). Sex differences in the effects of allopregnanolone on yohimbine-induced reinstatement of cocaine seeking in rats. Drug and Alcohol Dependence, 107(2–3), 264. https://doi.org/10.1016/j.drugalcdep.2009.11.002

Back, S., Dansky, B. S., Coffey, S. F., Saladin, M. E., Sonne, S., & Brady, K. T. (2000). Cocaine dependence with and without post-traumatic stress disorder: a comparison of substance use, trauma history and psychiatric comorbidity. The American journal on addictions, 9(1), 51–62. https://doi.org/10.1080/10550490050172227

Back, S. E., Brady, K. T., Jackson, J. L., Salstrom, S., & Zinzow, H. (2005). Gender differences in stress reactivity among cocaine-dependent individuals. Psychopharmacology, 180(1), 169–176. https://doi.org/10.1007/s00213-004-2129-7

Balopole, D. C., Hansult, C. D., & Dorph, D. (1979). Effect of cocaine on food intake in rats. Psychopharmacology, 64(1), 121–122. https://doi.org/10.1007/BF00427356

Bane, A. J., McCoy, J. G., Stump, B. S., & Avery, D. D. (1993). The effects of cocaine on dietary self-selection in female rats. Physiology & Behavior, 54(3), 509–513. https://doi.org/10.1016/0031-9384(93)90244-A

Becker, J. B., & Hu, M. (2008). Sex differences in drug abuse. Frontiers in Neuroendocrinology, 29(1), 36–47. https://doi.org/10.1016/j.yfrne.2007.07.003

Blain, L. M., Galovski, T. E., & Robinson, T. (2010). Gender differences in recovery from posttraumatic stress disorder: A critical review. Aggression and Violent Behavior, 15(6), 463–474. https://doi.org/10.1016/j.avb.2010.09.001

Brady, K., Grice, D., Dustan, L., & Randall, C. (1993). Gender Differences in Substance Use Disorders. Am J Psychiatry, 150(11), 1707–1711. https://doi.org/10.1176/ajp.150.11.1707

Brady, K., & Randall, C. (1999). Gender Differences in Substance Use Disorders. Psychiatric Clin North Am, 22(2), 241–52. https://doi.org/10.1016/S0193-953X(05)70074-5

Brodnik, Z. D., Black, E. M., Clark, M. J., Kornsey, K. N., Snyder, N. W., & España, R. A. (2017). Susceptibility to traumatic stress sensitizes the dopaminergic response to cocaine and increases motivation for cocaine. Neuropharmacology, 125, 295–307. https://doi.org/10.1016/j.neuropharm.2017.07.032

Bruening, A. B., Perez, M., & Ohrt, T. K. (2018). Exploring weight control as motivation for illicit stimulant use. Eating Behaviors, 30, 72–75. https://doi.org/10.1016/j.eatbeh.2018.06.002

Buffalari, D. M., Baldwin, C. K., Feltenstein, M. W., & See, R. E. (2012). Corticotrophin releasing factor (CRF) induced reinstatement of cocaine seeking in male and female rats. Physiology & Behavior, 105(2), 209–214. https://doi.org/10.1016/j.physbeh.2011.08.020

Buffalari, D. M., & See, R. E. (2009). Footshock stress potentiates cue-induced cocaine-seeking in an animal model of relapse. Physiology & Behavior, 98(5), 614–617. https://doi.org/10.1016/j.physbeh.2009.09.013

Buffalari, D. M., & See, R. E. (2011). Inactivation of the bed nucleus of the stria terminalis in an animal model of relapse: Effects on conditioned cue-induced reinstatement and its enhancement by yohimbine. Psychopharmacology, 213(1), 19–27. https://doi.org/10.1007/s00213-010-2008-3

Christiansen, D., Elklit, A., Christiansen, D., & Elklit, A. (2012). Sex Differences in PTSD. In Post Traumatic Stress Disorders in a Global Context. IntechOpen. https://doi.org/10.5772/28363

Cochrane, C., Malcolm, R., & Brewerton, T. (1998). The role of weight control as a motivation for cocaine abuse. Addictive Behaviors, 23(2), 201–207. https://doi.org/10.1016/S0306-4603(97)00046-4

Coffey, S. F., Saladin, M. E., Drobes, D. J., Brady, K. T., Dansky, B. S., & Kilpatrick, D. G. (2002). Trauma and substance cue reactivity in individuals with comorbid posttraumatic stress disorder and cocaine or alcohol dependence. Drug and Alcohol Dependence, 65(2), 115–127. https://doi.org/10.1016/S0376-8716(01)00157-0

Covington, H. E., Kikusui, T., Goodhue, J., Nikulina, E. M., Hammer, R. P., & Miczek, K. A. (2005). Brief Social Defeat Stress: Long Lasting Effects on Cocaine Taking During a Binge and Zif268 mRNA Expression in the Amygdala and Prefrontal Cortex. Neuropsychopharmacology, 30(2), Article 2. https://doi.org/10.1038/sj.npp.1300587

Covington, H. E., & Miczek, K. A. (2001). Repeated social-defeat stress, cocaine or morphine. Effects on behavioral sensitization and intravenous cocaine self-administration “binges.” Psychopharmacology, 158(4), 388–398. https://doi.org/10.1007/s002130100858

Doncheck, E. M., Liddiard, G. T., Konrath, C. D., Liu, X., Yu, L., Urbanik, L. A., Herbst, M. R., DeBaker, M. C., Raddatz, N., Van Newenhizen, E. C., Mathy, J., Gilmartin, M. R., Liu, Q., Hillard, C. J., & Mantsch, J. R. (2020). Sex, stress, and prefrontal cortex: Influence of biological sex on stress-promoted cocaine seeking. Neuropsychopharmacology, 45(12), 1974–1985. https://doi.org/10.1038/s41386-020-0674-3

Edwards, S., Baynes, B. B., Carmichael, C. Y., Zamora-Martinez, E. R., Barrus, M., Koob, G. F., & Gilpin, N. W. (2013). Traumatic stress reactivity promotes excessive alcohol drinking and alters the balance of prefrontal cortex-amygdala activity. Translational Psychiatry, 3(8), e296–e296. https://doi.org/10.1038/tp.2013.70

Erb, S., Shaham, Y., & Stewart, J. (1996). Stress reinstates cocaine-seeking behavior after prolonged extinction and a drug-free period. Psychopharmacology, 128(4), 408–412. https://doi.org/10.1007/s002130050150

Ersche, K. D., Stochl, J., Woodward, J. M., & Fletcher, P. C. (2013). The skinny on cocaine: Insights into eating behavior and body weight in cocaine-dependent men. Appetite, 71, 75–80. https://doi.org/10.1016/j.appet.2013.07.011

Evans, S. M., & Foltin, R. W. (2006). Exogenous Progesterone Attenuates the Subjective Effects of Smoked Cocaine in Women, but not in Men. Neuropsychopharmacology, 31(3), 659–674. https://doi.org/10.1038/sj.npp.1300887

Evans, S. M., Haney, M., & Foltin, R. W. (2002). The effects of smoked cocaine during the follicular and luteal phases of the menstrual cycle in women. Psychopharmacology, 159(4), 397–406. https://doi.org/10.1007/s00213-001-0944-7

Feltenstein, M. W., Henderson, A. R., & See, R. E. (2011). Enhancement of cue-induced reinstatement of cocaine-seeking in rats by yohimbine: Sex differences and the role of the estrous cycle. Psychopharmacology, 216(1), 53–62. https://doi.org/10.1007/s00213-011-2187-6

Fox, H. C., Sofuoglu, M., Morgan, P. T., Tuit, K. L., & Sinha, R. (2013). The effects of exogenous progesterone on drug craving and stress arousal in cocaine dependence: Impact of gender and cue type. Psychoneuroendocrinology, 38(9), 1532–1544. https://doi.org/10.1016/j.psyneuen.2012.12.022

Graf, E. N., Wheeler, R. A., Baker, D. A., Ebben, A. L., Hill, J. E., McReynolds, J. R., Robble, M. A., Vranjkovic, O., Wheeler, D. S., Mantsch, J. R., & Gasser, P. J. (2013). Corticosterone Acts in the Nucleus Accumbens to Enhance Dopamine Signaling and Potentiate Reinstatement of Cocaine Seeking. The Journal of Neuroscience, 33(29), 11800–11810. https://doi.org/10.1523/JNEUROSCI.1969-13.2013

Griffin, M. L., Weiss, R. D., Mirin, S. M., & Lange, U. (1989). A Comparison of Male and Female Cocaine Abusers. Archives of General Psychiatry, 46(2), 122–126. https://doi.org/10.1001/archpsyc.1989.01810020024005

Hadad, N. A., Wu, L., Hiller, H., Krause, E. G., Schwendt, M., & Knackstedt, L. A. (2016). Conditioned stress prevents cue-primed cocaine reinstatement only in stress-responsive rats. Stress, 19(4), 406–418. https://doi.org/10.1080/10253890.2016.1189898

Hegab, I., & Wei, W. (2014). Neuroendocrine changes upon exposure to predator odors. Physiology & Behavior, 131, 149–55. https://doi.org/10.1016/j.physbeh.2014.04.041

Hendrix, C. C., Anelli, L. M., Gibbs, J. P., & Fournier, D. G. (1994). Validation of the Purdue Post-Traumatic Stress Scale on a sample of Vietnam veterans. Journal of Traumatic Stress, 7(2), 311–318. https://doi.org/10.1007/BF02102951

Holly, E. N., Shimamoto, A., DeBold, J. F., & Miczek, K. A. (2012). Sex differences in behavioral and neural cross-sensitization and escalated cocaine taking as a result of episodic social defeat stress in rats. Psychopharmacology, 224(1), 179–188. https://doi.org/10.1007/s00213-012-2846-2

Institute of Laboratory Animal Resources, & National Research Council (Eds.). (2011). Guide for the care and use of laboratory animals (8th edn). National Acad. Press, 46(3). https://doi.org/10.1258/la.2012.150312

Jackson, L. R., Robinson, T. E., & Becker, J. B. (2006). Sex differences and hormonal influences on acquisition of cocaine self-administration in rats. Neuropsychopharmacology: Official Publication of the American College of Neuropsychopharmacology, 31(1), 129–138. https://doi.org/10.1038/sj.npp.1300778

Kessler, R. C., Sonnega, A., Bromet, E., Hughes, M., & Nelson, C. (1995). Posttraumatic Stress Disorder in the National Comorbidity Survey. Archives of General Psychiatry, 52(12), 1048. https://doi.org/10.1001/archpsyc.1995.03950240066012

Kondoh, K., Lu, Z., Ye, X., Olson, D. P., Lowell, B. B., & Buck, L. B. (2016). A specific area of olfactory cortex involved in stress hormone responses to predator odors. Nature, 532(7597), 103–106. https://doi.org/10.1038/nature17156

Kosten, T. R., Schottenfeld, R., Ziedonis, D., & Falcioni, J. (1993). Buprenorphine versus methadone maintenance for opioid dependence. The Journal of Nervous and Mental Disease, 181(6), 358–364. https://doi.org/10.1097/00005053-199306000-00004

Lacy, R. T., Austin, B. P., & Strickland, J. C. (2020). The influence of sex and estrous cyclicity on cocaine and remifentanil demand in rats. Addiction Biology, 25(1). https://doi.org/10.1111/adb.12716

Laque, A., Wagner, G. E., Matzeu, A., De Ness, G. L., Kerr, T. M., Carroll, A. M., Guglielmo, G., Nedelescu, H., Buczynski, M. W., Gregus, A. M., Jhou, T. C., Zorrilla, E. P., Martin-Fardon, R., Koya, E., Ritter, R. C., Weiss, F., & Suto, N. (2022). Linking drug and food addiction via compulsive appetite. British Journal of Pharmacology, 179(11), 2589–2609. https://doi.org/10.1111/bph.15797

Lynch, W. J., & Carroll, M. E. (1999). Sex differences in the acquisition of intravenously self-administered cocaine and heroin in rats. Psychopharmacology, 144(1), 77–82. https://doi.org/10.1007/s002130050979

Mantsch, J. R., Baker, D. A., Francis, D. M., Katz, E. S., Hoks, M. A., & Serge, J. P. (2007). Stressor- and corticotropin releasing factor-induced reinstatement and active stress-related behavioral responses are augmented following long-access cocaine self-administration by rats. Psychopharmacology, 195(4), 591–603. https://doi.org/10.1007/s00213-007-0950-5

McReynolds, J. R., Doncheck, E. M., Li, Y., Vranjkovic, O., Graf, E. N., Ogasawara, D., Cravatt, B. F., Baker, D. A., Liu, Q., Hillard, C. J., & Mantsch, J. R. (2018). Stress promotes drug seeking through glucocorticoid-dependent endocannabinoid mobilization in prelimbic cortex. Biological Psychiatry, 84(2), 85–94. https://doi.org/10.1016/j.biopsych.2017.09.024

Milivojevic, V., Fox, H. C., Sofuoglu, M., Covault, J., & Sinha, R. (2016). Effects of progesterone stimulated allopregnanolone on craving and stress response in cocaine dependent men and women. Psychoneuroendocrinology, 65, 44–53. https://doi.org/10.1016/j.psyneuen.2015.12.008

Moran-Santa Maria, M. M., Sherman, B. J., Brady, K. T., Baker, N. L., Hyer, J. M., Ferland, C., & McRae-Clark, A. L. (2018). Impact of endogenous progesterone on reactivity to yohimbine and cocaine cues in cocaine-dependent women. Pharmacology, Biochemistry, and Behavior, 165, 63–69. https://doi.org/10.1016/j.pbb.2017.11.001

Najavits, L. M., Gastfriend, D. R., Barber, J. P., Reif, S., Muenz, L. R., Blaine, J., Frank, A., Crits-Christoph, P., Thase, M., & Weiss, R. D. (1998). Cocaine Dependence With and Without PTSD Among Subjects in the National Institute on Drug Abuse Collaborative Cocaine Treatment Study. American Journal of Psychiatry, 155(2), 214–219. https://doi.org/10.1176/ajp.155.2.214

Najavits, L. M., Harned, M. S., Gallop, R. J., Butler, S. F., Barber, J. P., Thase, M. E., & Crits-Christoph, P. (2007). Six-Month Treatment Outcomes of Cocaine-Dependent Patients With and Without PTSD in a Multisite National Trial. Journal of Studies on Alcohol and Drugs, 68(3), 353–361. https://doi.org/10.15288/jsad.2007.68.353

Preston, K. L., & Epstein, D. H. (2011). Stress in the daily lives of cocaine and heroin users: Relationship to mood, craving, relapse triggers, and cocaine use. Psychopharmacology, 218(1), 29–37. https://doi.org/10.1007/s00213-011-2183-x

Preston, K. L., Kowalczyk, W. J., Phillips, K. A., Jobes, M. L., Vahabzadeh, M., Lin, J.-L., Mezghanni, M., & Epstein, D. H. (2018). Exacerbated Craving in the Presence of Stress and Drug Cues in Drug-Dependent Patients. Neuropsychopharmacology, 43(4), 859–867. https://doi.org/10.1038/npp.2017.275

Quadros, I. M. H., & Miczek, K. A. (2009). Two modes of intense cocaine bingeing: Increased persistence after social defeat stress and increased rate of intake due to extended access conditions in rats. Psychopharmacology, 206(1), 109–120. https://doi.org/10.1007/s00213-009-1584-6

Quiñones-Jenab, V., & Jenab, S. (2012). Influence of Sex Differences and Gonadal Hormones on Cocaine Addiction. ILAR Journal, 53(1), 14–22. https://doi.org/10.1093/ilar.53.1.14

Reed BG, Carr BR. The Normal Menstrual Cycle and the Control of Ovulation. [Updated 2018 Aug 5]. In: Feingold KR, Anawalt B, Blackman MR, et al., editors. Endotext [Internet]. South Dartmouth (MA): MDText.com, Inc.; 2000. https://www.ncbi.nlm.nih.gov/books/NBK279054/

Rogge, G., Jones, D., Hubert, G. W., Lin, Y., & Kuhar, M. J. (2008). CART peptides: Regulators of body weight, reward and other functions. Nature Reviews Neuroscience, 9(10), 747–758. https://doi.org/10.1038/nrn2493

Roth, M. E., & Carroll, M. E. (2004). Sex differences in the escalation of intravenous cocaine intake following long-or short-access to cocaine self-administration. Pharmacology Biochemistry and Behavior, 78(2), 199–207. https://doi.org/10.1016/j.pbb.2004.03.018

Saladin, M. E., Drobes, D. J., Coffey, S. F., Dansky, B. S., Brady, K. T., & Kilpatrick, D. G. (2003). PTSD symptom severity as a predictor of cue-elicited drug craving in victims of violent crime. Addictive Behaviors, 28(9), 1611–1629. https://doi.org/10.1016/j.addbeh.2003.08.037

Schreiber, A. L., Lu, Y. L., Baynes, B. B., Richardson, H. N., & Gilpin, N. W. (2017). Corticotropin-releasing factor in ventromedial prefrontal cortex mediates avoidance of a traumatic stress-paired context. Neuropharmacology, 113(Pt A), 323–330. https://doi.org/10.1016/j.neuropharm.2016.05.008

Schreiber, A. L., McGinn, M. A., Edwards, S., & Gilpin, N. W. (2019). Predator odor stress blunts alcohol conditioned aversion. Neuropharmacology, 144, 82–90. https://doi.org/10.1016/j.neuropharm.2018.10.019

Schwendt, M., Shallcross, J., Hadad, N. A., Namba, M. D., Hiller, H., Wu, L., Krause, E. G., & Knackstedt, L. A. (2018). A novel rat model of comorbid PTSD and addiction reveals intersections between stress susceptibility and enhanced cocaine seeking with a role for mGlu5 receptors. Translational Psychiatry, 8, 209. https://doi.org/10.1038/s41398-018-0265-9

Shin, K. M., Chang, H. Y., Cho, S.-M., Kim, N. H., Kim, K. A., & Chung, Y. K. (2015). Avoidance symptoms and delayed verbal memory are associated with post-traumatic stress symptoms in female victims of sexual violence. Journal of Affective Disorders, 184, 145–148. https://doi.org/10.1016/j.jad.2015.05.051

Sinha, R., Fox, H., Hong, K.-I., Sofuoglu, M., Morgan, P. T., & Bergquist, K. T. (2007). Sex steroid hormones, stress response, and drug craving in cocaine-dependent women: Implications for relapse susceptibility. Experimental and Clinical Psychopharmacology, 15(5), 445–452. https://doi.org/10.1037/1064-1297.15.5.445

Sofuoglu, M., Mitchell, E., & Kosten, T. (2004). Effects of progesterone treatment on cocaine responses in male and female cocaine users. Pharmacol Biochem Behav, 78(4), 699–705. https://doi.org/10.1016/j.pbb.2004.05.004

Somohano, V. C., Rehder, K. L., Dingle, T., Shank, T., & Bowen, S. (2019). PTSD symptom clusters and craving differs by primary drug of choice. Journal of Dual Diagnosis, 15(4), 233–242. https://doi.org/10.1080/15504263.2019.1637039

Substance Abuse and Mental Health Services Administration. (2017). Key substance use and mental health indicators in the United States: Results from the 2016 National Survey on Drug Use and Health. (HHS Publication No. SMA 17-5044, NSDUH Series H-52). Rockville, MD: Center for Behavioral Health Statistics and Quality, Substance Abuse and Mental Health Services Administration. Retrieved from https://www.samhsa.gov/data/

Substance Abuse and Mental Health Services Administration. (2021). Key substance use and mental health indicators in the United States: Results from the 2020 National Survey on Drug Use and Health. (HHS Publication No. PEP21-07-01-003, NSDUH Series H-56). Rockville, MD: Center for Behavioral Health Statistics and Quality, Substance Abuse and Mental Health Services Administration. Retrieved from https://www.samhsa.gov/data/

Takahashi, L. K., Nakashima, B. R., Hong, H., & Watanabe, K. (2005). The smell of danger: A behavioral and neural analysis of predator odor-induced fear. Neuroscience & Biobehavioral Reviews, 29(8), 1157– 1167. https://doi.org/10.1016/j.neubiorev.2005.04.008

Thanos, P. K., Cho, J., Kim, R., Michaelides, M., Primeaux, S., Bray, G., Wang, G.-J., & Volkow, N. D. (2011). Bromocriptine increased operant responding for high fat food but decreased chow intake in both obesity-prone and resistant rats. Behavioural Brain Research, 217(1), 165–170. https://doi.org/10.1016/j.bbr.2010.10.027

Tidey, J. W., & Miczek, K. A. (1997). Acquisition of cocaine self-administration after social stress: Role of accumbens dopamine. Psychopharmacology, 130(3), 203–212. https://doi.org/10.1007/s002130050230

Trappler, B., Braunstein, J. W., Moskowitz, G., & Friedman, S. (2002). Holocaust Survivors in a Primary Care Setting: Fifty Years Later. Psychological Reports, 91(2), 545–552. https://doi.org/10.2466/pr0.2002.91.2.545

Vicentic, A., & Jones, D. C. (2007). The CART (Cocaine- and Amphetamine-Regulated Transcript) System in Appetite and Drug Addiction. Journal of Pharmacology and Experimental Therapeutics, 320(2), 499– 506. https://doi.org/10.1124/jpet.105.091512

Wee, S., Specio, S. E., & Koob, G. F. (2007). Effects of Dose and Session Duration on Cocaine Self-Administration in Rats. Journal of Pharmacology and Experimental Therapeutics, 320(3), 1134–1143. https://doi.org/10.1124/jpet.106.113340

Weera, M. M., Schreiber, A. L., Avegno, E. M., & Gilpin, N. W. (2020). The role of central amygdala corticotropin-releasing factor in predator odor stress-induced avoidance behavior and escalated alcohol drinking in rats. Neuropharmacology, 166, 107979. https://doi.org/10.1016/j.neuropharm.2020.107979

Whitaker, A. M., & Gilpin, N. W. (2015). Blunted Hypothalamo-pituitary Adrenal Axis Response to Predator Odor Predicts High Stress Reactivity. Physiology & Behavior, 147, 16–22. https://doi.org/10.1016/j.physbeh.2015.03.033

Wortley, K. E., Chang, G.-Q., Davydova, Z., Fried, S. K., & Leibowitz, S. F. (2004). Cocaine- and amphetamine-regulated transcript in the arcuate nucleus stimulates lipid metabolism to control body fat accrual on a high-fat diet. Regulatory Peptides, 117(2), 89–99. https://doi.org/10.1016/j.regpep.2003.08.005

